# Dynamic competition between selective attention and spatial prediction during visual search

**DOI:** 10.1101/2025.07.10.664103

**Authors:** Floortje G. Bouwkamp, Jorie J.G. van Haren, Floris P. de Lange, Eelke Spaak

## Abstract

During visual search, we rely on both selective attention and spatial predictions to guide our behavior. However, whether and how these mechanisms interact is largely unclear. Using a contextual cueing paradigm, we investigated whether learning and exploitation of spatial predictive context can occur outside attentional focus. Repeating search scenes enabled distractor context to serve as a contextual cue predicting target location. Participants searched for a target among distractors in two colors: the same as the target-to-be-searched (attended context) or a different color (ignored context). Halfway through the experiments, we changed the target color, thereby altering the attentional status of distractor contexts while maintaining their spatial predictiveness. In Experiment 1, where participants regularly switched between target colors, we found exploitation of spatial predictive context both within and outside attentional focus. However, in Experiment 2, where attention was more stably focused on one target color, only the attended predictive context was exploited before transfer. Intriguingly, after transfer, previously ignored predictive context showed immediate benefits, revealing latent learning. These findings demonstrate a dynamic competition between selective attention and spatial predictions: while learning occurs independently of attention, exploitation may require attentional selection. Our results suggest that selective attention gates the influence of spatial predictions on behavior, with gating strength determined by the stability of attentional control.

**Significance statement:** This research addresses a fundamental question in cognitive science: can we learn from visual information we are not paying attention to? Our findings reveal a dynamic competition between selective attention and spatial prediction that can explain conflicting findings in the literature. We demonstrate that specifically the ability to exploit predictive spatial context depends critically on attentional stability. When required to flexibly switch attentional sets, we can benefit from both attended and unattended predictive information. However, when attention remains focused, unattended information is learned, but only becomes accessible when it is part of our attentional focus. These results reveal that the filtering operation of selective attention and the utilization of spatial prediction during visual search may actually compete dynamically, with selective attention ultimately constraining predictions’ influence on behavior. This advances our understanding of how these fundamental cognitive mechanisms interact under competition.

## Introduction

Our visual environment is usually crowded, making it impossible to process everything at once. Being able to locate a relevant item in such crowded scenes, something we refer to as visual search, is a ubiquitous skill needed for goal-driven perception in everyday life.

We know that selective attention is a crucial factor for efficient processing of our (visual) environment. By filtering out irrelevant items, we can devote cognitive resources to what matters for the goal at hand. For example, when searching for a phone, only items similar to the shape and color of the phone will be highlighted, speeding up the search process.

Additionally, we can make use of the predictable structure in our environment, as items tend to co-occur. These co-occurrence relations can generate predictions about *what* to expect and *where* to expect it (Chun & Turk-Browne, 2007; Peelen et al., 2023; Spaak et al., 2022; Theeuwes et al., 2022; Võ et al., 2019). To return to the previous example: when searching for a phone, specifically counter- and table tops are likely to be searched, as phones are most often found there. The detection of these types of regularities is generally thought to be implicit and the exploitation of such regularities a highly automatic process. This type of learning is called *visual statistical learning (VSL)*.

Within visual search, statistical learning of environmental regularities has been demonstrated to boost performance in several ways. For example, spatial regularities may cue the most likely location of a target, a.k.a *location probability cueing* (Geng & Behrmann, 2005; Geyer et al., 2024; Jiang et al., 2013), or set up expectations about where distracting information will be (Beesley et al., 2016; Ferrante et al., 2023; Richter et al., 2024). These types of cueing create a fixed attentional bias in space. Another type of cueing within visual search is *contextual cueing* (Chun & Jiang, 1998a). In this type of statistical learning, the *spatial context* within visual search scenes predicts where the target will be, thereby setting up an implicit spatial *relationship* (Jiang, 2018).

In a typical contextual cueing experiment, participants need to search for a target (often a letter T) that is embedded in a scene filled with similar-looking shapes (often rotated Ls). Some of these scenes are repeated over the course of the experiment, and although participants demonstrate little or no awareness of these repetitions (but see Meyen et al., 2024; Vadillo et al., 2016), they become markedly faster in finding the target in these repeated ‘old’ scenes compared to ‘new’ scenes. The consensus is that after learning the spatial relationship between distractors and target, this knowledge is exploited, which leads to more efficient guidance of attention when searching these scenes (Bouwkamp et al., 2025; Goujon et al., 2015; Jiang et al., 2019; Sisk et al., 2019; Spaak & de Lange, 2020). In other words, because the spatial context is predictive of target location, participants become more efficient at visual search. The underlying mechanisms of attention and prediction are unified in the concept of an attentional priority map. Bottom-up salience, goal-driven relevance, and prior experience together shape these priority maps (Duncan et al., 2023; Fecteau & Munoz, 2006; Ferrante et al., 2018; Sisk et al., 2019; Wolfe, 2021). These maps help us allocate attentional resources to whatever needs to be prioritized. But if, and how selective attention and predictions interact is largely unclear.

One big question that has occupied the field is how automatic visual statistical learning is. More specifically, is attention necessary for statistical learning to occur? When J.R. Saffran (1996) showed that children as young as 8 months old could extract regularities while passively listening to speech streams, this mechanism was thought to be highly automatic and attention was assumed to be unnecessary for this type of learning to occur. However, in the visual domain, it has been proposed that statistical learning, and contextual cueing specifically, though implicit, requires attentional selection (Jiang & Chun, 2001; Turk-Browne et al., 2005), limiting the scope of learning. If, instead, we could learn implicitly even from things we are ignoring, this would be incredibly useful, as it would be a less costly operation that could potentially run ‘in the background’. Moreover, the acquired knowledge might become relevant and thus beneficial at some future point in time. This attractive idea has led to multiple efforts to further investigate the role of selective attention in statistical learning. Both in the auditory and visual domain, statistical learning has been demonstrated to still occur when a secondary demanding task is requiring attention (Batterink & Paller, 2019; Musz et al., 2015), contradicting previous findings. Similarly, contextual cueing appears to be robust to distracting secondary tasks (Vicente-Conesa et al., 2022; Vickery et al., 2010).

A third stance is that selective attention is not required for statistical *learning* in visual search, but it is for the *exploitation* of what is learned. This has been suggested for contextual cueing when loading (visual) working memory with a secondary task (Goujon et al., 2015; Pollmann, 2019; Sisk et al., 2019), but also for contextual cueing of *ignored* visual context (Jiang & Leung, 2005). In the latter study, distractors in search scenes were present in two colors; however, the target was always in one and the same color. This narrowed down the search space as participants could safely ignore distractors in the non-target color (Wolfe, 2021). Contextual cueing did not occur for context that was being ignored, replicating the previous finding of Jiang & Chun (2001). However, when the task relevance of the distractor context was reversed by a change of color, there was a behavioral benefit when the previously ignored context was predictive. Jiang & Leung concluded that spatial predictive context was in fact learned, but that this learning was exploited only when the context became task-relevant and thus attended. This is thought to be evidence of ‘*latent learning’*. This latent learning phenomenon however failed to replicate recently (Vadillo et al., 2020), potentially calling into question its robustness.

In daily life, we change our *search goals* all the time (e.g., “Where is my phone?”, “Where is the traffic light?”, “Where is my friend?” etc.), while *contextual regularities* tend to stay stable over time (a room will not suddenly change color or shape). Though a feature change is inevitable in a transfer-paradigm, changing the color of the context might be more impactful than changing the color of the target. Moreover, for the association between context and target to become apparent after a change of context color, the association must be (latently) learned independently from its color feature.

Although it seems that mainly global configuration is learned, contextual cueing is sensitive to changes of perceptual identity of distractor items (Chun & Jiang, 1998a; Jiang & Wagner, 2004; Spaak & de Lange, 2020). If not only location but also color of distractor context is encoded during contextual learning (Jiang & Song, 2005), a color change of the context could be disruptive to the association between context and target, potentially erasing any contextual cueing.

In our current study, we investigated the role of selective attention on the learning and exploitation of spatial predictive context: can we learn from spatial context that is outside attentional focus? We used a contextual cueing paradigm very similar to the seminal studies by Jiang & Chun (2001) and Jiang & Leung (2005). The crucial difference is that after initial context learning, we applied a *target* color change in the transfer phase, while distractor context stayed exactly the same. Before this target color change we expected only spatial predictive context that is attended to be exploited. If, however, ignored spatial predictive context is latently learned, we expected to see the expression of this learning (or ‘exploitation’) to be uncovered directly after transfer when it becomes attended.

To foreshadow our results, we found the expected exploitation of predictive visual context that was attended. Surprisingly, we also found an effect of predictive visual context that could be ignored, already before transfer. Learning was therefore not ‘latent’, and its expression not dependent on attention. We reasoned that this exploitation of ignored spatial context may be due to a dynamic competition between selective attention and spatial predictions. Attention was regularly switching between colors due to alternating target color blocks, and this switching likely enabled spatial predictive context to influence behavior even from to-be-ignored locations. We confirmed this hypothesis in our follow-up experiment, where maintaining a consistent search goal (i.e., removing the target color switching) eliminated the exploitation of to-be-ignored predictive context before transfer. Interestingly, after transfer, this previously ignored context could immediately be exploited, suggesting that while learning occurs independently of attention, exploitation may require attentional selection. This pattern reveals a fundamental tension: while attentional filtering consistently reduces noise by restricting the search space, it may simultaneously block access to potentially beneficial predictive information. Together, these findings demonstrate that selective attention ultimately gates the influence of spatial predictions on behavior, with the strength of this gating is determined by the stability of attentional control.

*Transparency and Openness.* All data and code used for stimulus presentation and analysis are freely available on the Donders Repository at https://doi.org/10.34973/mqqe-jb24 This dataset has been subjected to a FAIR review protocol. *Note: This doi will become active upon publication*.

*Reviewers can access the data via a temporary link*.

We report how we determined our sample size, all data exclusions (if any), all manipulations, and all measures in the study. A priori power analyses using G*Power were used to calculate sample sizes. These studies were not preregistered.

## Experiment 1

## Material and methods

### Participants

An a priori calculation determined that to reliably detect a small effect size (Vadillo et al. 2020, experiment 3) with 80% power at alpha level .05, we required a sample size of N=104. A total of 109 participants were^1^ recruited in 2021 via the prolific platform (http://www.prolific.co) to participate in an online experiment. All participants had normal or corrected to normal vision, normal hearing and no history of neurological or psychiatric conditions. They provided written informed consent and received financial reimbursement for their participation in the experiment. The study followed the guidelines for ethical treatment of research participants by CMO 2014/ 288 region Arnhem-Nijmegen, The Netherlands. Participants that performed with an accuracy below 50% during any of the blocks were automatically excluded from the dataset. Furthermore we excluded seven participants with an overall accuracy score that fell below the 25^th^ percentile minus 1.5 x interquartile range. This resulted in a final sample size of 102 participants (Age: 31.3±6.5 years, 35 males^1^) with an average performance of 94.87% (*SD*= 3.14) on the task. All participants gave written informed consent beforehand and were paid for their participation. To motivate participants, an overall performance level >90% correct was rewarded with an additional bonus pay-out.

### Stimuli & Apparatus

Search scenes consisted of 17 stimuli, 1 target letter T and 16 L-shaped distractors, measuring 1.2° × 1.2° in size. Distractors L shapes were rotated with a random multiple of 90°and had a 10% offset in the line junction to increase search difficulty (Y. Jiang & Chun, 2001). Stimuli were placed on a regular grid spanning from −9° to +9°in 9 steps horizontal and −6° to +6° in 6 steps vertical from the center of the screen. Random jitter of ±0.4° was added to stimuli locations to prevent collinearities with other stimuli (Chun & Jiang, 1998a). To control difficulty of the scenes, the target always appeared between 6° and 8° of eccentricity, and the mean distance between the target and all distractors was kept between 9° and 11°. The target was tilted either to the left (−90°) or to the right (+90°). Stimuli were presented using the Gorilla platform (https://gorilla.sc/) and custom-written JavaScript. Participants performed the task online on their own device (tablet and phone excluded). Size of stimuli was controlled with a standard visual degree check implemented in Gorilla and the task could only be done in full screen mode.

### Procedure

Participants were instructed to find a target letter T that was hidden amongst L-shaped distractors, and report how this target was tilted with a key press on their keyboard; the left ‘C’ key for tilted leftward and the right ‘M’ key for tilted rightward (Figure 1). Participants searched for a black target for an entire block, after which they had to search for a white target for an entire block, and these blocks kept alternating until the end of the experiment. The target color of the first block was randomized across participants. Half of the distractors were the same color as the target, the other half of the distractors were in the other color, and all were presented on a grey background (see also experimental design). To remind people of the upcoming target color, each block started with the target letter T shown upright in the correct color, serving as a cue (2.5 s). Throughout the experiment there was a fixation dot presented at the center of the screen in same color as the target, as a continuous reminder of the current target color. Participants were asked to fixate on this dot between search trials, but were allowed to freely move their eyes when searching. Each trial started with a brief fixation period (1 sec), then search scenes where displayed until response or up to 3 s, after which the response on trial was registered as too late. After this, participants received feedback on the trial for 500 ms: the fixation dot was replaced with a green ‘+’ indicating the answer was correct, a red ‘x’ when the response was incorrect, and a blue ‘o’ if they were too late. The experiment started with task instructions and a gorilla screen calibration, followed by the main task consisting of 2 practice blocks (one white target color block and one black target color block) and 28 experimental blocks (14 white target color blocks and 14 black target color blocks). After each block participants received feedback on their performance, indicating average response speed and % correct of that block, as well as overall % correct as they would be rewarded if the latter would be above 90%. Participants could take a short break after each block. At the end of the experiment there was a short questionnaire on their experience.

**Figure 1.**
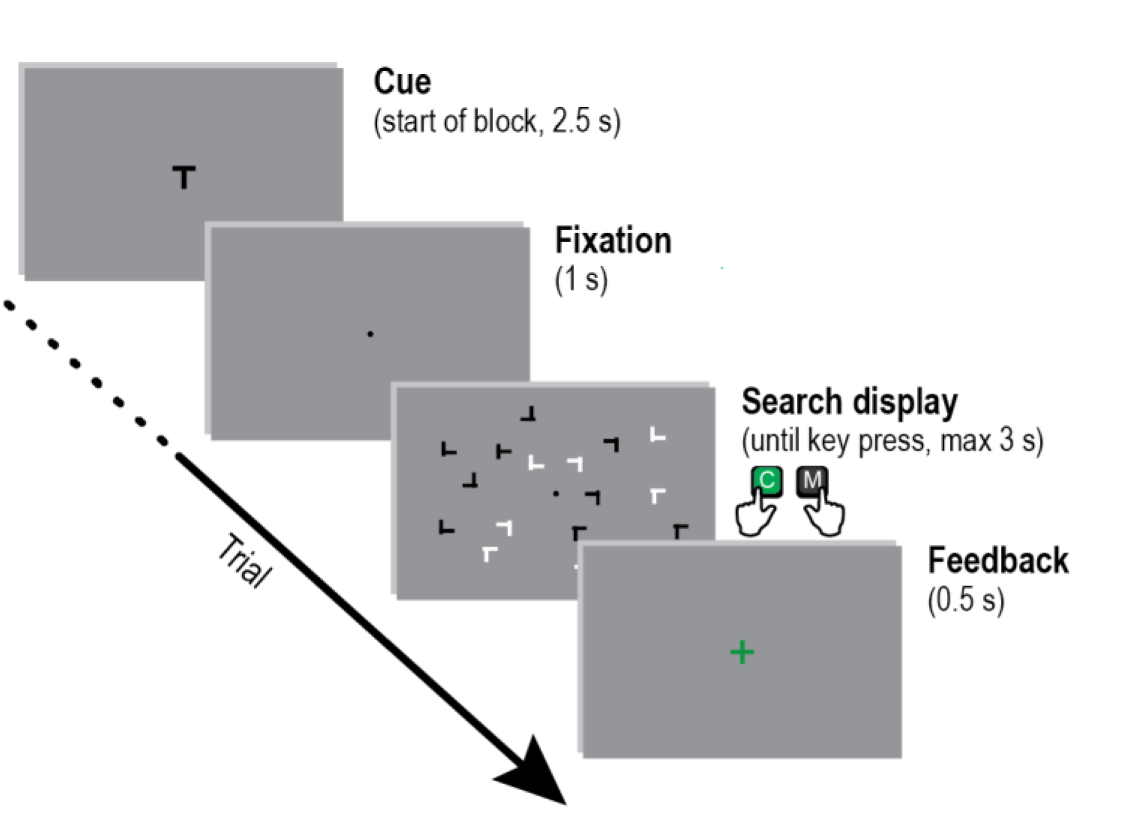
Task. Participants performed a visual search task where they had to locate a target letter T embedded amongst distractor L-shapes and report its orientation with a left (‘C’) or right (‘M’) key press. A block started with a target letter T cue in the color assigned to that block. Feedback was given at the end of each trial (correct, incorrect, or too late). All figures use example scenes with a black target color.

### Experimental design

Participants searched for either a black or a white target, depending on the block. On all trials half of the distractors were white, the other half were black. This resulted in one distractor set that was the same color as the target, and one set which was of a different color. As it is typically assumed that participants can limit their (serial) search to one attended color, we label the target-color-matching distractors the *attended context*. The other distractor set we label the *ignored context* (Figure 2A). We manipulated predictability by repeating or not repeating distractor context along with the target location. If spatial context was *predictive* in a scene, both the location and orientation of the distractor set was repeated in subsequent blocks (‘old’ in contextual cueing parlance). The target location was also repeated if spatial context was predictive, but the target orientation was always randomized, to prevent motor response learning. If not repeated in subsequent blocks, the locations of the distractor set would be generated randomly each time, making them ‘new’ and thus *nonpredictive* (Figure 2B). These two factors, attentional status and predictiveness, were independently manipulated. Combined, they led to all scenes having *attended context*, that could be *predictive* (ATT+) or *nonpredictive* (ATT-), and *ignored context* that could be *predictive* (IGN+) or *nonpredictive* (IGN-). This generated four conditions (Figure 2B): both the attended and the ignored spatial context was predictive, labeled as *all context predictive* (ALL CP = ATT+IGN+), only the attended spatial context was predictive, labeled as *attended context predictive* (ATT CP = ATT+IGN-), only the ignored spatial context was predictive, labeled as *ignored context predictive* (IGN CP = ATT-IGN+), or neither the attended, nor the ignored spatial context was predictive, labeled as *no context predictive* (NO CP = ATT-IGN-).

**Figure 2.**
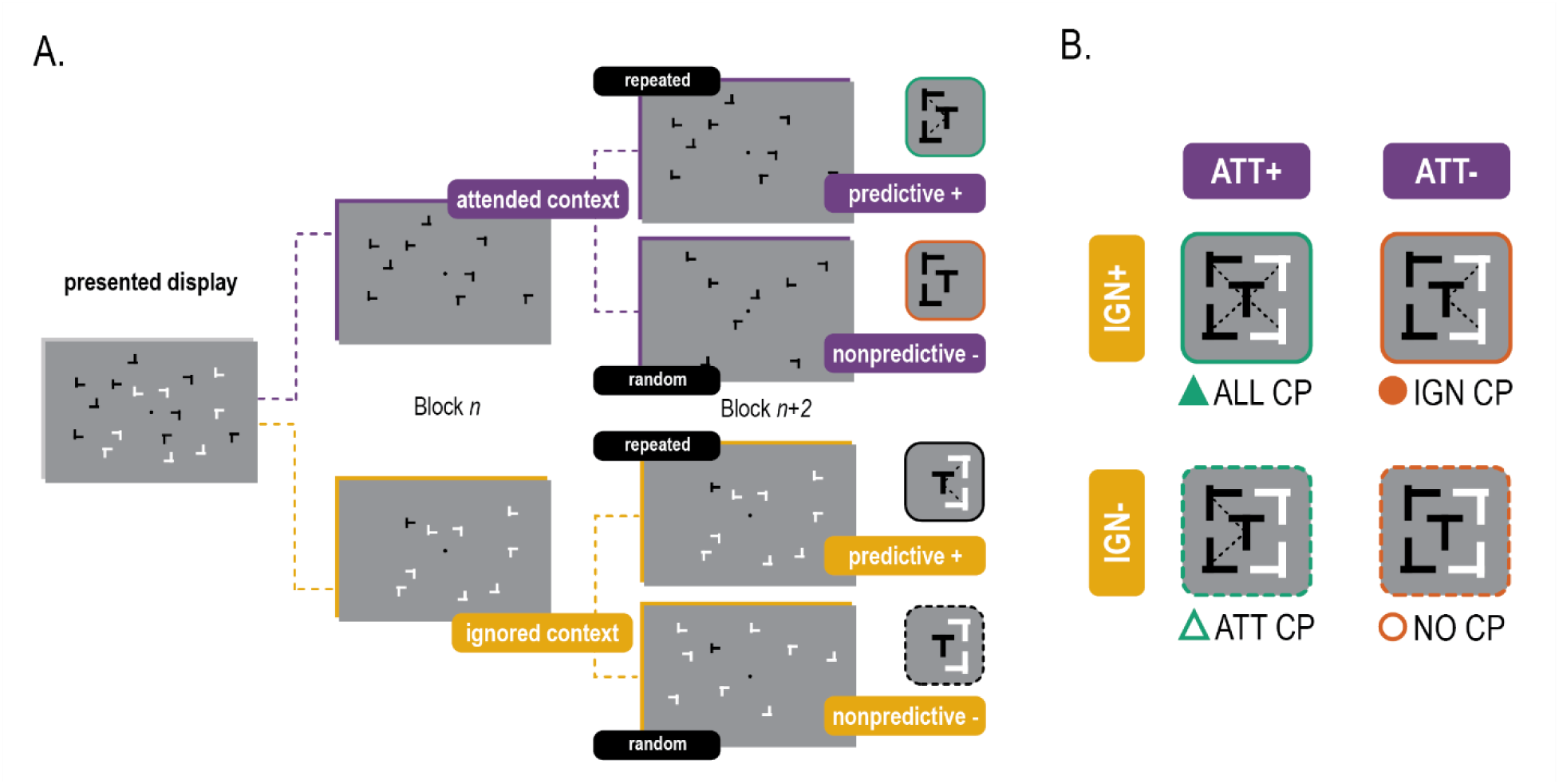
Experimental manipulations. **A)** The color of the distractor context determined the attentional status. In this example the target is black, therefore black distractor context is attended and white distractor context is ignored. Furthermore, the location and orientation of the distractors was either repeated over blocks or created randomly. This determined the predictiveness of the distractor context, reflected additionally in the pictograms on the right side: when connected with dashed lines to the T, the context is predictive**. B)** Attentional status (ATT or IGN) and Predictiveness (+/-) combined generated our four conditions: all context predictive (ALL CP), attended context (ATT CP) predictive, ignored context predictive (IGN CP) and no context predictive (NO CP).

Crucially, approximately half way through the experiment we reversed the attentional status of the spatial context by changing the target color, a moment we call *transfer* (Figure 3). In doing so the spatial context remains identical but task relevance swaps: the distractor set that was previously attended now became ignored, and the distractor set that was ignored became attended (Figure 3A). Because participants were always searching black and white targets in alternating blocks, this change remained entirely implicit. The transfer has no effect on the NO CP condition as there is no predictiveness in the scenes (ATT-IGN-→ IGN-ATT-). Both contexts in the ALL CP condition stay predictive after the transfer, but the attended predictive context becomes ignored, and the ignored predictive context becomes attended (ATT+IGN+ → IGN+ATT+). The ATT CP condition and IGN CP condition essentially trade places due to the transfer: scenes where the attended context was predictive (ATT CP) change into scenes where the ignored context is predictive (ATT+IGN-→ IGN+ATT-). The reverse is true for scenes where the ignored context was predictive (IGN CP: ATT-IGN+ → IGN-ATT+), they becomes scenes where the attended context is predictive (Figure 3B).

**Figure 3.**
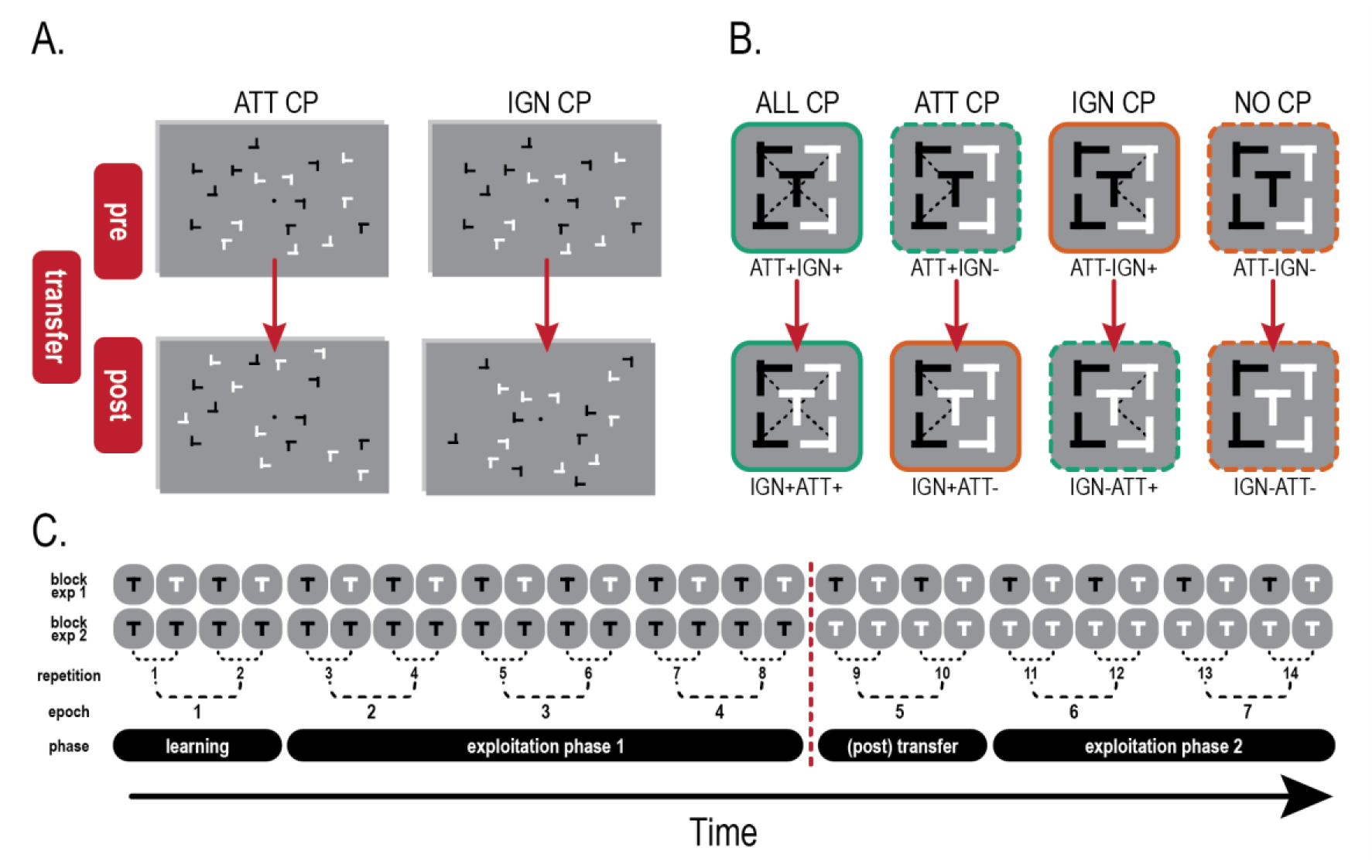
The transfer. **A)** Example of the transfer for two of our four conditions, and only for when the target letter T was black. Left is a scene where the attended context is predictive of target location. After transfer the black distractor context is exactly the same and thus repeated, however, the target color changed from black to white. Effectively, the predictive context becomes ignored. On the right an example where the predictive context was ignored, and the target color change at transfer had the opposite effect. **B)** Transfer for all four conditions depicted in pictograms. **C)** At the top the search goal structure over the time course of the experiment is depicted: for experiment 1 this was alternating per block, for experiment 2 this was consistent until transfer. Below the aggregation scheme for our different phases used for analyses. Transfer occurred after 4 out of 7 epochs, indicated with the red dotted line.

Finally, we mimicked ‘stay’ and ‘switch’ trials from previous literature (Jiang & Chun, 2001; Vadillo et al., 2020) by adding a fifth condition: another all context predictive condition, but without a transfer (ALL CP-nt: ATT+IGN+ → ATT+IGN+). Each of the 28 blocks consisted of 4 scenes per condition with a target in each quadrant of the visual field to prevent target probability learning (Jiang et al., 2013). This adds up to 20 trials per target color block, and in total 40 unique scenes, 8 per condition. These scenes were repeated (or created newly in the NO CP condition) for 14 times in total, so every scene was repeated every second block (hence ‘repetition’ in figure 3C). The transfer was applied after 8 repetitions, leaving 6 repetitions for ‘new’ learning. As commonly done, we averaged over two repetitions, generating 7 epochs (consisting of 16 trials per condition) and these were then assigned to four phases. During the initial *learning phase* we expect no exploitation of spatial predictive context as a minimum of two repetitions is needed to learn (Tseng & Lleras, 2013). Subsequently this learning can continue, but can also be visible as an behavioral advantage, which we label as *the exploitation phase 1*. The epoch directly after transfer is the crucial *(post) transfer phase* where we can see how the transfer impacts exploitation behavior. And lastly, to assess exploitation of newly acquired spatial predictive context we have the final *exploitation phase 2* (figure 3C).

### Data Analysis

Data were both analyzed and visualized using R (R Core Team, n.d.) using packages *ggplot2* (Wickham, 2016), *ggrain* (Allen et al., 2021), *ez* (Lawrence, n.d.).Reaction time was our primary, and accuracy our secondary dependent variable of interest. Only trials with a response given within the time limit were included (98.18% of trials), and trials with an response given before 400 msec were regarded as accidental presses and excluded (5 trials). Reaction time analysis was performed on correct responses only. We first assessed whether our experimental manipulation was successful with a 2×2 repeated measures ANOVA with time (all 7 epochs) and condition (all 5 levels) as factors, expecting both main effects and an interaction. We then analyzed the data in a priori defined phases of the experiment: after initial learning (epoch 1, 2 repetitions), the first exploitation phase started (epoch 2-3-4). Between epoch 4 and 5 we applied the transfer and switched the target color in the scenes. The epoch directly after this is called the post transfer phase, and this where we expect to see the effect of the transfer. Lastly, after there has been an opportunity for new learning, there is the second exploitation phase (epoch 6 and 7). The exploitation phases (1 and 2) were analyzed with a 2×2 repeated measures ANOVA with Attended context (predictive versus nonpredicitive) and Ignored context (predictive versus nonpredicitive) as factors. The impact of transfer was assessed by adding transfer (pre/post) as a factor, resulting in a 2×2×2 repeated measures ANOVA. We additionally contrasted conditions both pre and post transfer using two sided t-tests with a Holm-Bonferroni correction for multiple comparisons (planned contrasts), which is the same as the interaction between conditional difference and transfer. This allowed us to test the effect of transfer while removing the general task improvement over time, assuming this to be equal across conditions. The ALL CP-nt (no transfer) condition was analyzed separately. First, we hypothesized that if changing the attentional status of spatial predictive context altered exploitation of the ALL CP condition, performance in the ALL CP condition should be worse compared to the ALL CP-nt (no transfer) condition, but only post transfer. We tested this with a one-sided t test on the difference between the ALL CP conditions (transfer/no transfer) pre versus post transfer. Secondly, we explored how a difference, if any, between these conditions post transfer, subsequently behaved over time with a 2×2 ANOVA, with factors condition (ALL CP and ALL CP-nt) and epoch (5-6-7).

For our ANOVA’s, if Mauchly’s test was significant, and thus the assumption of sphericity was violated, we report the corrected p-value and degrees of freedom. Only when the Greenhouse–Geisser ε values were above 0.75 did we report the more liberal Huyn–Feldt corrected values (Field et al., 2012). For F-tests, the generalized eta-squared measure of effect size (η^2^G) is reported (Bakeman, 2005). For T-tests we report Cohen’s d as effect size, using the approach of Gibbons et al., (n.d.) for paired samples, including a suggested correction by Borenstein (2009).

## Results

Participants searched for either a black or a white target, depending on the block. Half of all distractors shared the color with the target (the *attended* distractor set), while the other half had a different color (the *ignored* distractor set). Additionally, distractor context was either repeated across blocks (*predictive*) or created randomly (*nonpredictive*).

### Accuracy

Overall accuracy was near ceiling levels, indicating participants performed well on the visual search task (96.62 ± 2.41). There were no significant differences between white target and black target blocks (*t*_101_ = .26, *p*= .792, *d*=.015). Importantly, although the accuracy of participants improved over time (*F*_3.7,373_ = 20.02, *p*_GG_ <.001, η^2^G= .048), there were no condition differences in the accuracy rates (*F*_3,303_ = 1.83, *p*= .142, η^2^G= .001), nor the improvement over time (*F*_13.5,1363_ = 1.33, *p*_GG_=.184, η^2^G= .006).

### Reaction times

#### General

Average response speed was well beneath the time limit (1421 ± 537 ms). There was no significant difference in response speed between black target and white target blocks (*t*_101_= 1.663, *p*= .099, *d*= .069). Participants improved over time, becoming faster at finding and reporting target orientation (*F*_4.6,468_ = 105.73, *p*_GG_ <.001, η^2^G= .138). There was a difference between our experimental conditions (*F*_2.89,292_ = 7.10, *p*_HF_ <.001, η^2^G= .005), and the improvement over time differed per condition (*F*_16.9,1703_ = 1.87, *p_HF_* =.014, η^2^G= .004). Our experimental manipulation thus affected response speed (Figure 4).

**Figure 4.**
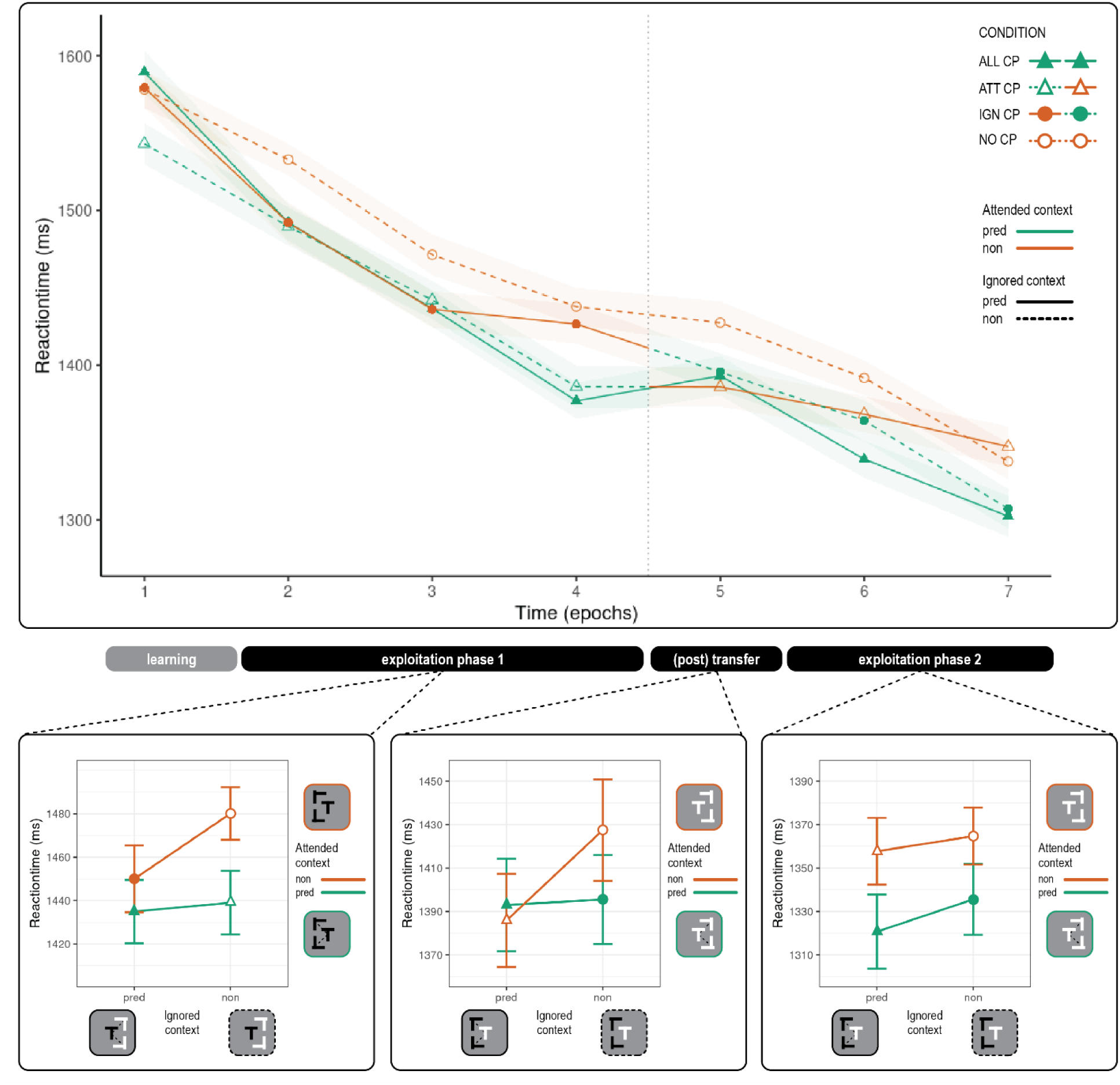
Results experiment 1. Top: Reaction time per condition plotted over the time course of the experiment. Shaded areas represent within subject corrected 95% confidence interval. Vertical dotted line between epoch 4 and 5 indicates target color change at transfer. Note how the line colors change after transfer, indicating a change of attentional status. Three panels below reflect the interaction analyses of reaction time per phase. From left to right: Exploitation phase1, post transfer, exploitation phase 2. Predictive status of ignored context is plotted on x-axis, predictive status of attended context is plotted in the two different colors. Note that for post-transfer and exploitation phase 2, the then-current (i.e., after transfer) predictive status is used to label conditions. Bars indicate within subject corrected 95% confidence interval.

#### Exploitation phase 1

We examined whether participants, after the initial learning blocks, could exploit the predictive context, for both the attended and ignored context. As expected, participants successfully exploited the attended spatial predictive context (*F*_1,101_= 10.32, *p* =.002, η^2^G= .006; *predictive* 1434 ± 530 ms vs *nonpredictive* 1461 ± 543 ms, Figure 4 & 5), confirming previous findings. However, in contrast to previous reports, and to our surprise, participants could also exploit the *ignored* spatial predictive context (*F*_1,101_= 4.07, *p* =.046, η^2^G= .002; *predictive* 1439 ± 532 vs *nonpredictive* 1456 ± 541 ms, Figure 4 & 5). There was no interaction between the effects of attended and ignored context (*F*_1,101_= 2.67, *p* =.105, η^2^G= .001).

**Figure 5.**
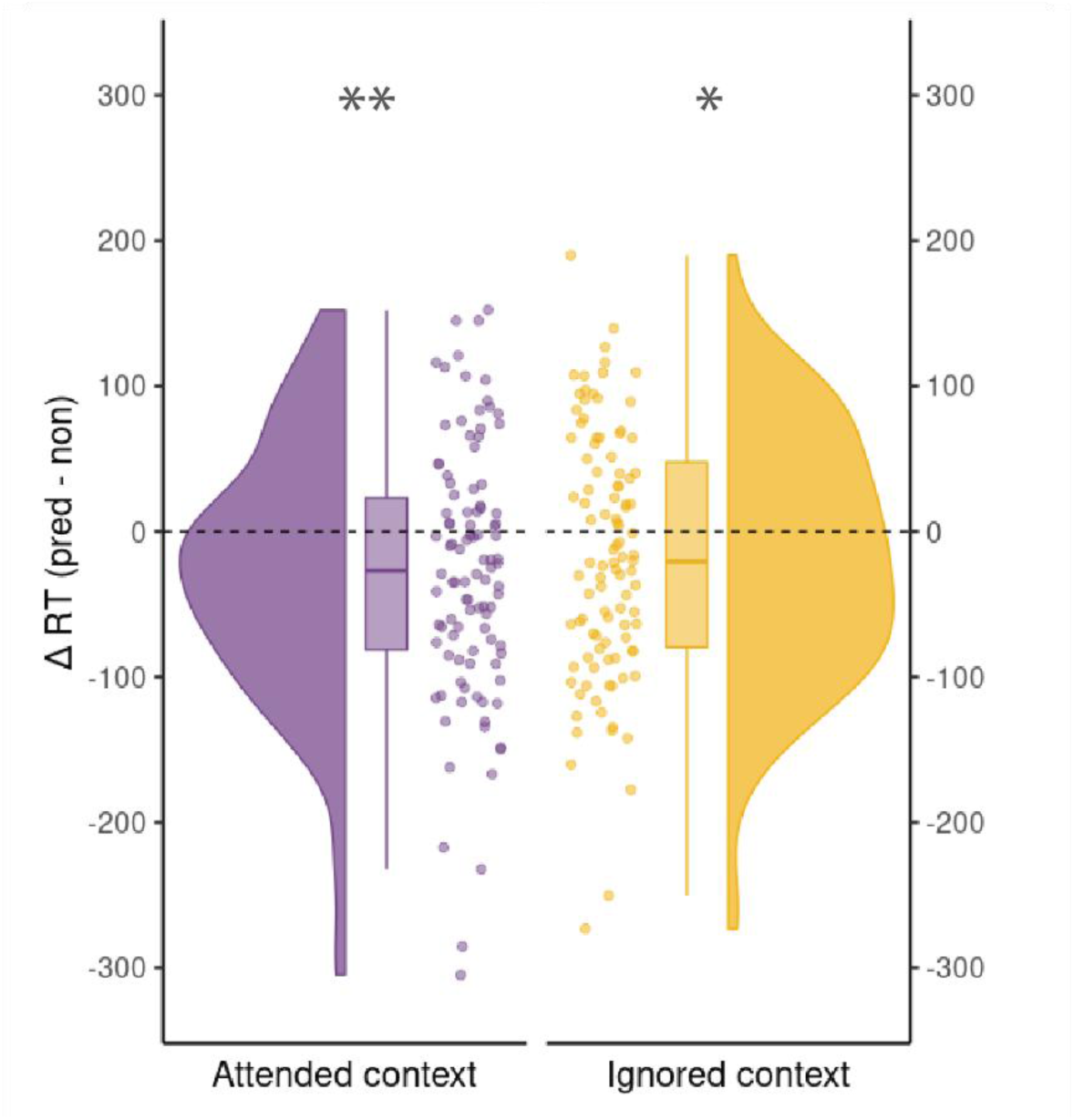
Distribution of the main effect (predictive versus nonpredictive) for both Attended context (left in purple) and Ignored context (right, in yellow) of experiment 1, exploitation phase 1. Negative values represent faster reaction times when the context is predictive versus when it is not.

#### Pre/Post Transfer before and after target color change

Little over half way through the experiment, we introduced the target color transfer: scenes that had a white target color before, now had a black target and scenes that had a black target before, now had a white target. The rest, i.e. the repetitions and the color of the distractor contexts, remained the same. This effectively changed the attended context to ignored, and the ignored spatial context to attended. When comparing the first exploitation phase (epoch 2-4) to search behavior directly after the transfer (epoch 5), we see that both the exploitation of the spatial predictive context that was attended before transfer, and the exploitation of spatial predictive context that was ignored before transfer, persisted, even though the target color changed (Attended context: *F*_1,101_= 6.63, *p*=.015, η^2^G= .003; Ignored context:: *F*_1,101_= 5.71, *p*=.019, η^2^G= .003). Adding the post transfer phase additionally revealed an interaction between the effect of Attended context and Ignored context (*F*_1,101_= 4.22, *p*=.042, η^2^G= .002). Across phases, Ignored predictive context was exploited specifically when the Attended context was *not* predictive of target location (IGN CP condition). As expected, there is a general increase in performance over time (*F*_1,101_= 27.00, *p*< .001, η^2^G= .018), but none of the patterns changed from the exploitation phase to post transfer (no interaction with phase pre/post, all *p*-values >.05). We thus find no evidence of a sudden change in exploitative behavior, neither in the negative (loss of advantage) nor positive direction (gain of advantage). Our planned contrasts support these results: none of the differences between conditions before the transfer was impacted by the target color change (all *p* values >.05). Importantly, the exploitation of spatial predictive context that was ignored (but in our case, exploited nonetheless), did not improve significantly when it became attended due to the target color change at transfer. We thus find no evidence that pre transfer learning leads to a sudden exploitation when spatial predictive context becomes task-relevant after transfer, a.k.a. ‘latent learning’. This is perhaps not surprising, considering we found, in contrast to previous reports, that ignored spatial predictive context is already exploited before transfer. Taken together, we can conclude that spatial context that is predictive of target location is exploited, both before and after target color change at transfer. Does that mean the transfer had no impact at all? We can directly compare what happens when *both* attended and ignored predictive context are swapped due to the transfer (ALL CP) to when there is no swap at all (ALL CP-nt; both contexts predictive but no color swap at transfer, see Methods for details). We tested whether the transfer negatively impacted performance, and this is indeed what we find (*t*_101_ = 1.70, *p*=0.046, *d*= 0.22). After transfer this difference remained for the rest of the experiment (time: *F*_2,202_= 25.03, *p*< .001, η^2^G= .032; transfer/no transfer: *F*_1,101_= 5.13, *p=* .026, η^2^G= .005; no interaction: *F*_2,202_=.89, *p=*.413, η^2^G< .001). While the former results demonstrate that spatial predictive relations are learned and exploited independently of overall attention, this comparison indicates that the attentional set under which the relations were learned is nonetheless a relevant feature.

#### Exploitation phase 2

After post transfer exposure to the new target color, we found that participants learned to exploit the ‘new’ spatial predictive relations only for the attended context (*F*_1,101_= 14.67, *p*< .001, η^2^G= .010). After transfer we no longer see evidence of the exploitation of spatial predictive context that is ignored (Ignored context: *F*_1,101_= 1.47, *p=* .228, η^2^G= .001; no interaction *F*_1,101_=.16, *p*= .686, η^2^G< .001). We thus conclude that both spatial predictive context that is attended *and* that is ignored can be exploited during visual search, and this exploitative behavior was neither suddenly nor drastically impacted when the target color changes at transfer, making the previously ignored spatial context attended and vice versa. Instead, we see a slow increase in the exploitation of previously ignored spatial predictive context when it becomes attended and a slow decrease of exploitation of previously attended spatial predictive context when it becomes ignored.

## Discussion

The fact that we see exploitation of predictive spatial context in ignored-color distractors specifically in the initial exploitation phase is in contrast with previous findings (Y. Jiang & Chun, 2001; Vadillo et al., 2020). We wondered whether this transient nature was due to the switching between white and black targets, which occurred after every block of trials. It could be that the requirement to switch between black target color and white target color blocks generated a more ‘open’ attentional filter compared to when participants would have to solely focus on one target color all the time. This would also explain why the target color change at transfer was less impactful than expected. Moreover, if people not only improve in the task in general, but also improve in the task of ‘focusing on the correct color’, this would explain why we do *not* see this exploitation of ignored spatial predictive context in the latter part of the experiment. Put differently, our results may indicate a tug of war between the filtering operation of attention and the exploitation of spatial predictive context in the task-irrelevant color.

To test this hypothesis, we executed a follow up experiment, which was identical to the first experiment, except for one critical difference: participants no longer alternated in searching black and white targets throughout the experiment. Instead, they searched for one target color until transfer. Then, at transfer, they were given explicit instructions that the target color was changed. If the alternating nature of the search during the first experiment resulted in an ‘open’ attentional filter, and therefore to the exploitation of spatial predictive context even when it is ignored, this effect should be absent in this non-alternating version of the experiment.

## Experiment 2

## Material & Methods

### Participants

All procedures were exactly the same as the first experiment. We recruited 108 participants in 2024 via the prolific platform (http://www.prolific.co) to participate in the online experiment. All participants had normal or corrected to normal vision, normal hearing and no history of neurological or psychiatric conditions. They provided written informed consent and received financial reimbursement for their participation in the experiment. The study followed the guidelines for ethical treatment of research participants by CMO 2014/ 288 region Arnhem-Nijmegen, The Netherlands. To ensure people were naïve, participation in our first experiment was an exclusion criterion in the recruitment for the second experiment. Six participants with an overall accuracy score that fell below the 25^th^ percentile minus the 1.5 x interquartile range, were excluded. This resulted in a final sample size of 102 participants (Age: 30.3±7.5 years, 50 males) with an average performance of 93.94% (*SD*= 3.52%) on the task.

### Procedure

The procedure was identical to the first experiment except for one aspect: participants did not search for white and black targets alternating in blocks. Instead, they searched for a target of one color (black target: N=51, white target: N=51) until transfer, when the target color changed. Participants received short instructions at the transfer point indicating that from now on the target color was changed.

### Experimental design

As in Experiment 1, forty unique scenes were generated, 8 per condition, but this time all with the same target color. At transfer the target color changed, changing the attentional status of the distractor sets, except for the ALL CP-nt condition. As we were now forced to change the target color also in this condition, we additionally changed the color of the distractor context. For these scenes, since *both* target and distractor context change color, predictive context that is attended stays attended and the predictive context that is ignored, stays ignored.

### Data Analysis

We excluded trials that were too late (2.22 %) and trials with a response time under 400 ms (6 trials). After cleaning the data, we used one and the same pipeline for data analyses of both the first and second experiment.

## Results

### Accuracy

Overall accuracy on the target orientation task was again near ceiling (96.06 ± 2.72%). There was no difference between participants who started with a white color target and those who started with a black colored target (*t*_98.347_ = 1.12, *p*= 0.264, *d*= .22). Mirroring the results of experiment 1, participants improved over time (*F*_3.7,373_ = 20.02, *p*_GG_ <.001, η^2^G= .048), but conditions did not differ in accuracy (*F*_3,303_ = 0.33, *p*= .801, η^2^G< .001), nor did the improvement over time depend on condition (*F*_16.5,1669_ = .95, *p*_HF_=.517, η^2^G= .004).

Reaction times

#### General

Average response speed was highly similar to experiment 1 (1436 ± 545 ms). There was no difference between participants that started with a black target compared to those that started with a white target (*t*_98,685_= 0.36, *p*= 0.717, *d*= .07). Participants improved over time, becoming faster at finding and reporting target orientation (*F*_4.5,452_ = 96.99, *p*_GG_ <.001, η^2^G= .115). There was a difference between our experimental conditions (*F*_3,303_ = 3.90, *p*= .009, η^2^G= .003), and the improvement over time also depended on condition (*F*_17.5,1765_ = 1.86, *p*_HF_= .016, η^2^G= .004). Again, our experimental manipulation had an effect on response speed of the task (Figure 6, top).

**Figure 6.**
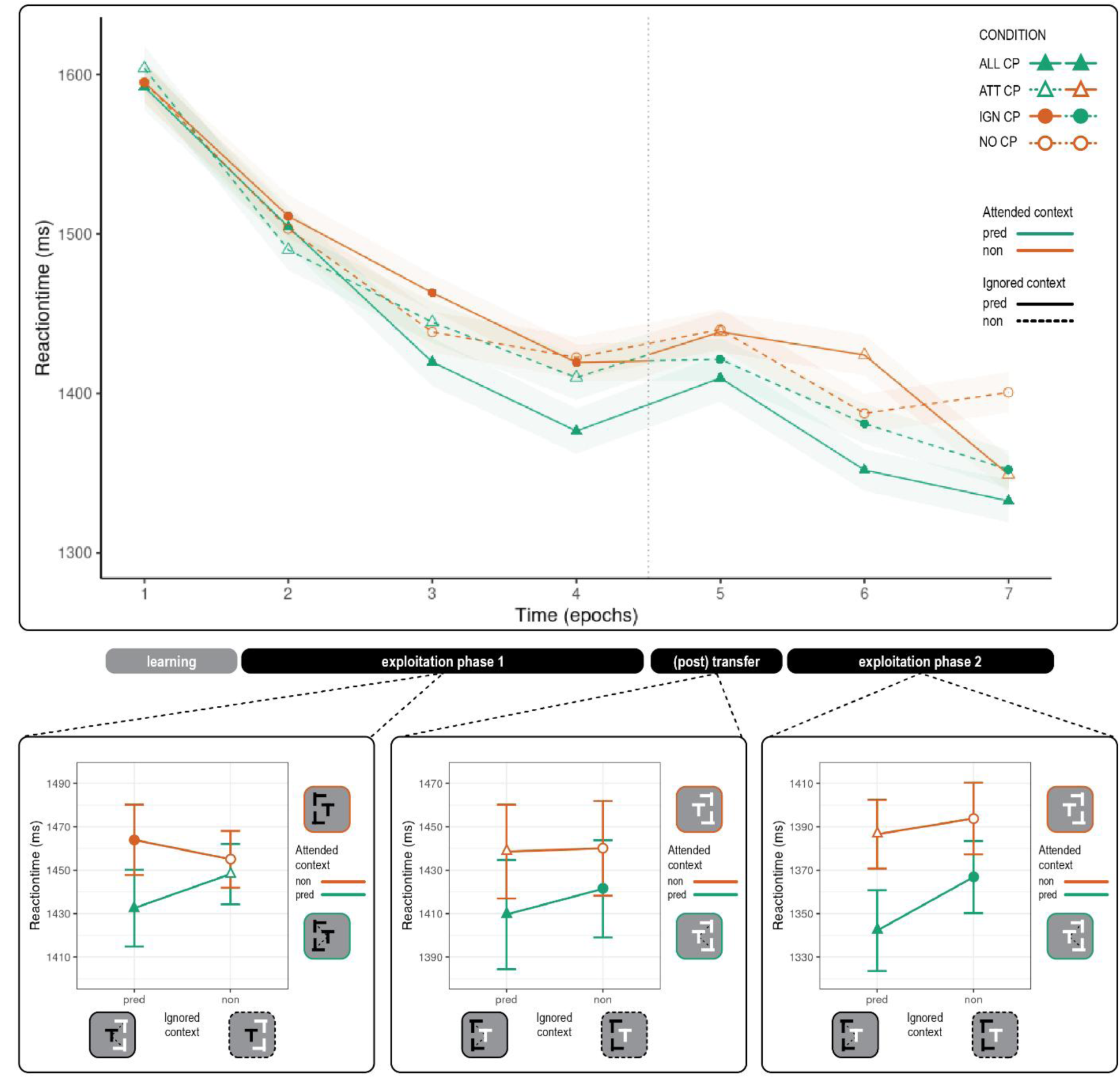
Results experiment 2. Top: Reaction time per condition plotted over the time course of the experiment. Shaded areas represent within subject corrected 95% confidence interval. Vertical dotted line between epoch 4 and 5 indicates target color change at transfer. Note how the line colors change after transfer, indicating a change of attentional status. Three panels below reflect the interaction analyses of reaction time per phase. From left to right: Exploitation phase 1, post transfer, exploitation phase 2. Predictive status of ignored context is plotted on x-axis, predictive status of attended context is plotted in the two different colors. Bars indicate within subject corrected 95% confidence interval.

#### Exploitation phase 1

After initial learning, participants exploit spatial predictive context that is attended (*F*_1,101_ = 6.96, *p*= .010, η^2^G= .003, *predictive* 1436 ± 543 ms vs *nonpredictive* 1456 ± 543 ms, Figure 6 & 7), similar to what we found in experiment 1. However, this time we find no evidence of exploitation of spatial predictive context that was being ignored (*F*_1,101_ = .13, *p*= .72, η^2^G< .001, *predictive* 1444 ± 543 ms vs *nonpredictive* 1447 ± 543 ms, Figure 6 & 7). We thus find, as hypothesized, that when target color search is stable, allowing for a more efficient filtering of color, people no longer use spatial predictive context in the ignored color. The question that remains is whether this context is also not latently learned.

**Figure 7.**
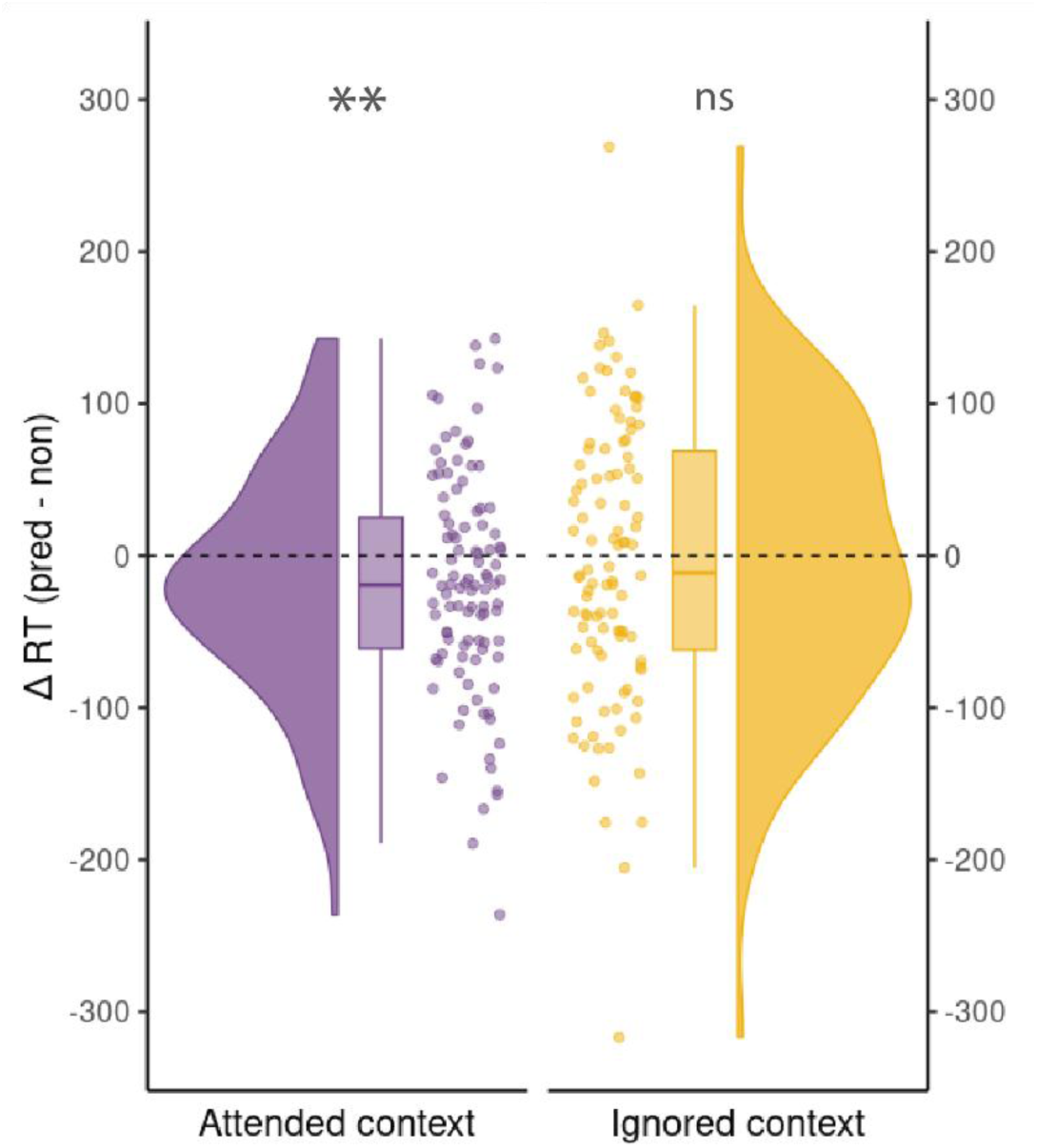
Distribution of the main effect for both Attended context (left in purple) and Ignored context (right, in yellow) of experiment 2, exploitation phase 1. Negative values represent faster reaction times when the context is predictive versus when it is not.

#### Pre/Post Transfer: before and after target color change

If only spatial predictive context that is attended can be learned and exploited, we should see the behavioral advantage of spatial predictive context that is attended disappear from pre to post transfer phase, as it can no longer be exploited now that it is ignored. If, however, spatial predictive context that was ignored before transfer was latently learned but requires attention to be exploited, we should see a behavioral advantage after transfer of previously ignored, but now attended spatial predictive context. This is, in fact, what we find. Besides a general improvement over time (*F*_1,101_ = 4.55, *p*= .03, η^2^G= .003) we find an effect of exploitation of spatial predictive context that was attended, *both* pre and post transfer (*F*_1,101_ = 8.12, *p*= .005, η^2^G= .003). We again, find no evidence of exploitation of spatial predictive context that was ignored (main effect *Ignored* context: *F*_1,101_ = .41, *p*= .521, η^2^G< .001; interaction *Attended x Ignored* context: *F*_1,101_ = .83, *p*= .364, η^2^G< .001), and none of the patterns changed from the exploitation phase to post transfer (no interaction with phase pre/post, all *p*-values >.05). These results indicate that immediately after the target color change at transfer, spatial predictive context that is attended is exploited. Importantly, this spatial predictive context was still ignored pre-transfer. This would imply this spatial predictive context was, in fact, *latently* learned, but not exploited, during the initial exploitation phase. Instead this learning is uncovered when the spatial predictive context became attended post-transfer, leading to exploitation. Similar to our first experiment, the target color change at transfer had little impact on the exploitation of either predictive spatial context that was attended and now ignored or vice versa: all planned contrasts on conditional differences pre versus post transfer were non-significant. Contrary to our findings in our first experiment, we also find no (negative) impact of transfer when directly comparing the all context predictive conditions (*transfer < no transfer*, *t*_101_= 1.55, *p*= .062, *d*= .20). Participants still improved behaviorally in both ALL CP conditions during the latter part of the experiment (*F*_2,202_ = 19.59, *p*< .001, η^2^G< .023), but the lack of a difference between *transfer* and *no transfer* persisted (*F*_1,101_ = 1.51, *p*= .222, η^2^G= .002) and did not depend on time (*F*_2,202_ = .36, *p*= .695, η^2^G< .001).

#### Exploitation phase 2

After the opportunity to learn after transfer, we see the that spatial predictive context is exploited, but only when this spatial predictive context is attended (*F*_1,101_ = 13.01, *p*< .001, η^2^G= .009) and not when it was ignored (*F*_1,101_ = 2.64, *p*= .107, η^2^G= .002), and there was also no interaction (*F*_1,101_ = .74, *p*= .392, η^2^G< .001).

## Discussion

These results show that when the task allows for a stable attentional filter, spatial predictive context that was ignored can no longer be exploited. We do see, however, *post transfer* exploitation of spatial predictive context that was ignored *pre transfer*, when it becomes attended due to the target color change. This is evidence of latent learning of ignored spatial predictive context and in line with the findings of Jiang & Leung (2005). Our conclusion is that while learning spatial predictive seems to be possible independently of selective attention, the expression of that learning, a.k.a *exploitation,* of spatial predictive context *is* dependent on selective attention.

Our first experiment demonstrates that flexible attention enables both learning and exploitation of spatial predictive context, regardless of whether that context falls within or outside the attentional focus. Our second experiment reveals that while learning remains intact, the exploitation of these spatial regularities critically depends on an ‘open’ attentional filter: eliminating task switching (as present in Experiment 1) abolishes the exploitation of predictive spatial context in the ignored color, with learned regularities only benefiting performance once they become attended. This pattern reveals a dynamic competition between selective attention and spatial predictions: when attention is flexible due to task switching, spatial predictions can influence behavior even from ignored locations, but when attentional selection is stable and efficient, it gates the exploitation of predictive information. Together, these experiments illuminate how selective attention and spatial predictions engage in a complex interplay during visual search, with their relative influence determined by the stability of attentional control.

## General Discussion

Here, we investigated whether learning and exploitation of spatial predictive context in visual search can occur without attention. Levering a *contextual cueing* paradigm, we generated spatial predictions during visual search: repeated distractor context becomes predictive of target location and this enhances attentional guidance through these scenes (Chun, 2000; Chun & Jiang, 1998b; Goujon et al., 2015; Sisk et al., 2019). Additionally we altered the attentional status of parts of the scene by manipulating their color. Distractors of the same color as the to-be-searched target need to be inspected, and are therefore attended. Conversely, distractors in the other color could be ignored. Crucially, approximately halfway through the experiments, we changed the color of the target (transfer) within the scenes, changing the attentional status of the distractor contexts. Combining spatial predictiveness and attentional status enabled us to investigate how goal-directed selective attention and predictions interact. As expected, we found robust learning and exploitation of spatial predictive context that was task-relevant and therefore attended. Surprisingly, and counter to earlier work (Jiang & Chun, 2001; Jiang & Leung, 2005), we also observed both learning *and* exploitation of *ignored* spatial predictive context. The exploitation of ignored spatial context however only occurred when participants regularly switched in terms of the color of the searched-for target, potentially leading to a broader attentional filter. When participants’ attention was more stable (Experiment 2), participants still learnt but no longer exploited the ignored spatial context. This latent learning became visible when the spatial context became attended after transfer. We discuss these findings below.

As expected, spatial predictive context that was task-relevant and thus attended was learned and exploited: in both experiments participants quickly learned and were able to exploit attended spatial context that was predictive of target location to improve visual search performance. This adds to a large body of evidence that indicates that when spatial predictive context is attended, it can be learned and exploited (Chun, 2000; Chun & Turk-Browne, 2007; Goujon et al., 2015; Jiang & Chun, 2001; Sisk et al., 2019). However, in our first experiment, we additionally found learning and exploitation of spatial predictive context that was task-irrelevant and thus could be ignored.

Importantly, we found this effect already before the change of attentional status at transfer. Learning from ignored spatial predictive context was therefore not latent but manifest: exploitation was not dependent on selective attention. Already in the classic study by Chun & Jiang (1998) evidence of the exploitation of ignored predictive context was found. However, when they increased the difficulty in the task by making the distractors more similar to the target, this effect disappeared. This finding was used to argue that when the task was too easy, attentional resources automatically spilled over to the irrelevant context. This was confirmed in a later study (Jiang & Leung, 2005). However, in this study they found that when the attentional status of ignored spatial predictive context changed, this context suddenly could be exploited. This was taken as evidence that the spatial relations in the ignored context were *latently* learned. In an extensive and high-powered study by Vadillo et al. (2020), this latent learning was not replicated. Vadillo et al. did, however, find evidence of the exploitation of ignored predictive context in one of their experiments, and argued this is due to participants inability to truly ignore the context in the task-irrelevant color. This dovetails with our findings. After observing the results from our first experiment, we hypothesized that the switching between target colors might have demanded more flexible attention, and as a consequence, created a more open attentional filter. This would lead to less efficient suppression of task irrelevant context, giving rise to an opportunity to learn and exploit this context when it was predictive of target location.

To investigate the influence of the flexibility of attention on the exploitation of spatial predictive context, we removed the task switching and instead gave participants a stable goal in a follow-up experiment. As expected, learning and exploitation from the attended spatial predictive context was unaffected while we no longer found evidence of exploitation of ignored spatial predictive context. We did however, find evidence for exploitation of ignored spatial predictive context when a transfer in task relevance changed its attentional status to attended. This suggests that this spatial predictive context was *latently* learned when it was still ignored, but required attention to be exploited. Taken together, these experiments yield two important insights. Firstly, selective attention is crucial for *exploitation* of spatial predictive context in visual search. People consistently learn and exploit spatial predictive context that is task relevant and thus attended. Moreover, if the irrelevant spatial context is efficiently filtered out, people can no longer exploit this context, even though it is predictive of target location. Secondly, *learning* spatial regularities is not dependent on selective attention. This is consistent with the early findings by Jiang & Leung (2005), but not with more recent work (Vadillo et al., 2020). One important distinction was that the change of attentional status of the spatial context at transfer was instantiated via a target color change, in both our experiments. This was a deliberate deviation from previous work investigating the role of selective attention and contextual learning (Jiang & Leung, 2005; Vadillo et al., 2020, 2024). What is typically done, is changing the color of all distractor stimuli at transfer. While this approach maintains consistent search goals and minimizes explicit awareness of the change, it may introduce unintended consequences. Specifically, the spatial context-target association is consistently paired with color and thus likely encoded with color information (Turk-Browne et al., 2008). Altering the distractor colors could disrupt the learned associations more severely than a simple target color change. Our results support this notion.

Instead of losing all behavioral advantage, as is commonly seen, the change at transfer was less impactful in both our experiments. For our first experiment, it can be argued that post transfer exploitation can be attributed to the open attentional filter that explains our results before transfer. However, this is less likely in our second experiment with stable and efficient selective attentional filtering. We believe our target color change allowed for more exploitation of the learned spatial context-target location association compared to changing the color of all distractors and this revealed latent learning. This is therefore an important consideration for future work on this topic.

The concept of an attentional filter stems from the well-established finding that selective attention enables people to restrict their visual search to a specific color-defined subset of elements. (Egeth et al., 1984; Kaptein et al., 1995; Palmer, 1994). The efficiency of visual search depends on how well selective attention can restrict processing: according to the Guided Search model (Wolfe, 2021) selective attention uses feature-based signals to guide the deployment of the spatial attentional ‘spotlight’. In our design that would mean that the target-defining feature of color creates priority maps through which the attentional spotlight has to move. A recent study by Duecker et al. (2024) investigated the difference between guided (i.e., color-cued) and unguided search of scenes consisting of stimuli in two colors. They found that guided search behavioral performance was indistinguishable from unguided search of half the set size. This reduction of the search space was accompanied by neural enhancement of the task relevant context and neural suppression of the task irrelevant context. We had a very similar set-up (amount of trials, stimuli and blocked target color cue), yet do not see such efficient suppression of the to be ignored spatial context. An important distinction is that in our experiments the ignored spatial context was a potential cue that could be learned and exploited. Interestingly, a recent study by Vadillo et al. (2024) found that the set size of the irrelevant context impacts visual search performance, demonstrating that this context was not perfectly ignored. However, this set size effect did not depend on attentional resources and did not interact with the contextual cueing effect: more attention to the irrelevant spatial context did not lead to more learning nor exploitation when it was predictive of target location. More research is necessary to fully understand when and how selective attention enables learning from spatial regularities.

An important consideration is to what degree the irrelevant predictive context could truly be ignored. As relevant and irrelevant context were spatially overlapping and the target could be anywhere, the irrelevant context could only be ignored via the mechanism of feature based selective attention. Recently Duncan et al. (2024) demonstrated that limiting the search space via an explicit color cue abolished the learning of the spatial distribution of distractors in the to-be-ignored color. In their design, however, color-based ignoring prevented further processing of the shapes, and with it access to predictive information, as it was a certain shape that occurred more often at one location. In our visual search scenes, suppression of distractors in one color automatically encompasses brief processing of the spatial context in that color, and it is this spatial context that can be predictive of target location. Moreover, in subsequent search both contexts are still encountered, as they are overlapping. In other lines of research, what we define as search space is referred to as the attentional window. The size of this window can be altered, for instance by using more central compared to peripheral stimuli (Kim & Jeong, 2025) and importantly, this can alter the impact of distracting stimuli. Cast in this perspective, our notion of poor filtering would translate to a larger attentional window as both would lead to more context being considered for search. It would be interesting to investigate if spatial predictive context that should *not* be considered as part of the search space by some other cue, can still impact search performance. For instance, marking the area where the target can be found would render the context outside of this area truly irrelevant.

Previous research has shown that delineating the search space this way, and with it set the attentional window, can modulate the impact of distractors (Biggs & Gibson, 2018). If one would still find an effect of context outside of the attentional window, this would run counter to the known local effect in contextual cueing, i.e. that mostly the context directly surrounding the target is responsible for enhanced search performance (Bouwkamp et al., 2025; Brady & Chun, 2007). However, this local effect has only been established for attended predictive context, without any filtering operation of selective attention that is global of nature. This is therefore an exciting avenue for future research on this topic.

The exploitation of spatial predictive context in ignored locations appears to follow a dynamic pattern across our experiments. In Experiment 1, this exploitation was primarily observed before transfer, with later phases showing exploitation only for attended contexts. This pattern suggests that attentional filtering mechanisms strengthen over time, eventually preventing the utilization of predictive information outside the attentional focus, despite its initial benefit to performance. This effect was even more pronounced in Experiment 2, where optimized filtering completely eliminated the exploitation of ignored predictive context until it became attended. These findings reveal a fundamental tension between spatial predictions and selective attention: while perfect filtering of task-irrelevant context optimizes processing efficiency, it simultaneously blocks access to potentially beneficial predictive information. This creates an interesting paradox where context that is predictive of target location could be considered ‘task-relevant’ by definition, yet attending to it requires the processing of more distractors than strictly necessary based on target color alone. While previous research has suggested that selective attention and predictions jointly modulate priority maps in the brain (Fecteau & Munoz, 2006; Ferrante et al., 2018; Sisk et al., 2019; Wolfe, 2021), our results demonstrate that these mechanisms may actually compete dynamically, with selective attention ultimately constraining prediction’s influence on behavior. This competition can be understood through a signal-to-noise framework: attentional filtering consistently reduces noise by restricting the search space, whereas the benefits of processing ignored distractors only materialize when they contain predictive information. This asymmetry in reliability may explain why attention ultimately gates the influence of spatial predictions, advancing our understanding of how these fundamental cognitive mechanisms interact under competition.

### Constraints on Generality

This study made use of large sample sizes and online recruitment enabled a diverse sample. This likely implies a greater generalization of our findings to the general population, compared to typical laboratory experiments conducted on small samples of undergraduate students. Our use of simplified visual search task might limit applicability to real world experience. While naturally occurring searches can be complex, these simplified paradigms nonetheless capture key aspects of real-world visual search (Botch et al., 2023).

---

**Supplementary figure 1.**
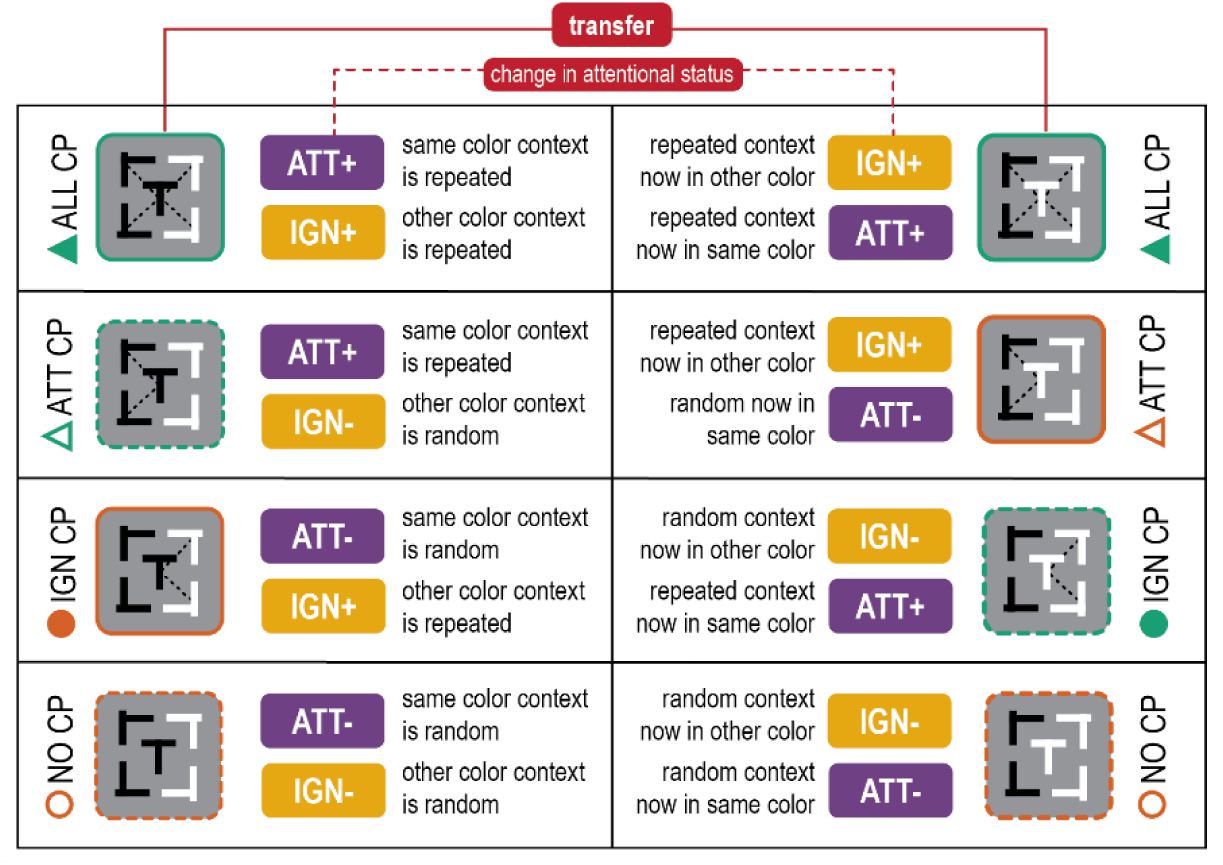
Conditions and transfer. Overview of all conditions labels, the attentional status and predictive status of the distractor context of scenes, together with a verbalization what that means at the scene level. In this instance the target was black pre-transfer. In experiment 1 there was another set of scenes with a white target pre-transfer. In experiment 2 the starting target color was counterbalanced across participants. Note how we kept condition names the same post-transfer, but change line type color and symbols. This is to clarify these scenes are the same scenes, the transfer impacts the attentional status of its context(s).

## Footnotes

1 Demographics of 98 of 102 participants, demographics of 4 participants were unknown

